# Speed and strength of an epidemic intervention

**DOI:** 10.1101/2020.03.02.974048

**Authors:** Jonathan Dushoff, Sang Woo Park

## Abstract

An epidemic can be characterized by its speed (i.e., the exponential growth rate *r*) and strength (i.e., the reproductive number ℛ). Disease modelers have historically placed much more emphasis on strength, in part because the effectiveness of an intervention strategy is typically evaluated on this scale. Here, we develop a mathematical framework for this classic, strength-based paradigm and show that there is a corresponding speed-based paradigm which can provide complementary insights. In particular, we note that *r* = 0 is a threshold for disease spread, just like ℛ = 1, and show that we can measure the speed and strength of an intervention on the same scale as the speed and strength of an epidemic, respectively. We argue that, just as the strength-based paradigm provides the clearest insight into certain questions, the speed-based paradigm provides the clearest view in other cases. As an example, we show that evaluating the prospects of “test-and-treat” interventions against the human immunodeficiency virus (HIV) can be done more clearly on the speed than strength scale, given uncertainty in the proportion of HIV spread that happens early in the course of infection. We suggest that disease modelers should avoid over-emphasizing the reproductive number at the expense of the exponential growth rate, but instead look at these as complementary measures.

## 1 Introduction

An epidemic can be described by its *speed* and *strength*. The speed of an epidemic is characterized by the exponential growth rate *r*, which measures how *fast* an epidemic grows at the population level. The strength of an epidemic is characterized by the reproductive number ℛ, which measures how many new cases are caused by a typical *individual* case. Knowing the speed and strength of an epidemic allows predictions about the course of the epidemic and the effectiveness of intervention strategies.

Much research has prioritized estimates of ℛ, and particularly its value in a fully susceptible population, called the *basic* reproductive number ℛ_0_, because ℛ has a threshold value (i.e., ℛ = 1) that determines whether a disease can invade, the level of equilibrium, and the effectiveness of control efforts (Anderson and May, 1991; Diekmann et al., 1990). The insight that a case must on average cause at least one new case under good conditions for a disease to persist goes back > 100 years (Ross, 1911); the idea of averaging by defining a ‘typical’ case was formalized 30 years ago (Diekmann et al., 1990). ℛ is also of interest because it provides a *prima facie* prediction about the total *size* of an epidemic (Anderson and May, 1991; Ma and Earn, 2006; Arino et al., 2007; Andreasen, 2011; Miller, 2012).

Here, we show that *r* can also serve as a threshold, and also provide a useful metric for difficulty of elimination. We first generalize the idea that ℛ measures the difficulty of elimination by showing we can measure an intervention’s “strength” on the same scale as the reproductive number. We then show that we can likewise measure an intervention’s “speed”, and that there is a duality between the threshold ℛ = 1 and a corresponding minimal intervention strength required for elimination, and the threshold *r* = 0 and a corresponding minimal intervention speed. We argue that the historical primacy of ℛ over *r* is partly artificial, and discuss cases where strength provides the better framing for practical disease questions and cases where speed does.

## 2 Methods

### 2.1 Epidemic model

We model disease incidence using the renewal equation, a simple, flexible framework that can cover a wide range of model structures (Heesterbeek and Dietz, 1996; Diekmann and Heesterbeek, 2000; Roberts, 2004; Aldis and Roberts, 2005; Wallinga and Lipsitch, 2007; Roberts and Heesterbeek, 2007; Champredon et al., 2018). In our model, disease incidence at time *t* is given by:

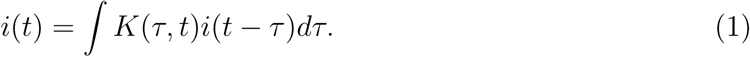

Here, *K*(*τ, t*) is the infection kernel describing how infectious we expect an individual infected *τ* time units ago to be in the population. In general, *K*(*τ, t*) will depend on population characteristics that may change through time *t* – notably, the proportion of the population susceptible, *S*(*t*). Since we are interested in invasion and control, we will generally neglect changes in *K*(*τ, t*) through time. Thus, we will assume *K*(*τ, t*) ≡ *K*(*τ*). Importantly, this means we are neglecting changes in susceptible proportion through time: *S*(*t*) ≈ *S*(0). Under this assumption, the renewal equation is equivalent to the Von Foerster equations (see e.g. Fraser et al. (2004)).

### 2.2 Strength-based decomposition

Assuming that the infection kernel *K* doesn’t change with time, we write:

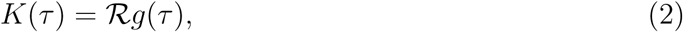

where *g*(*τ*) is the “intrinsic” generation-interval distribution. The generation interval is defined as the time between when a person becomes infected and when that person infects another person (Svensson, 2007); therefore, the intrinsic generation-interval distribution *g*(*τ*) gives the relative infectiousness of an average individual as a function of time since infection (Champredon and Dushoff, 2015). Since *g* is a distribution, it integrates to 1, and the basic reproductive ℛ number is thus the integral of *K*.

Imagine a control measure that proportionally reduces *K*, for example, by protecting a fixed fraction of susceptibles through vaccination (Fig. 1A). We then have:

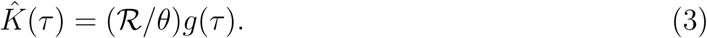

**Figure 1:**
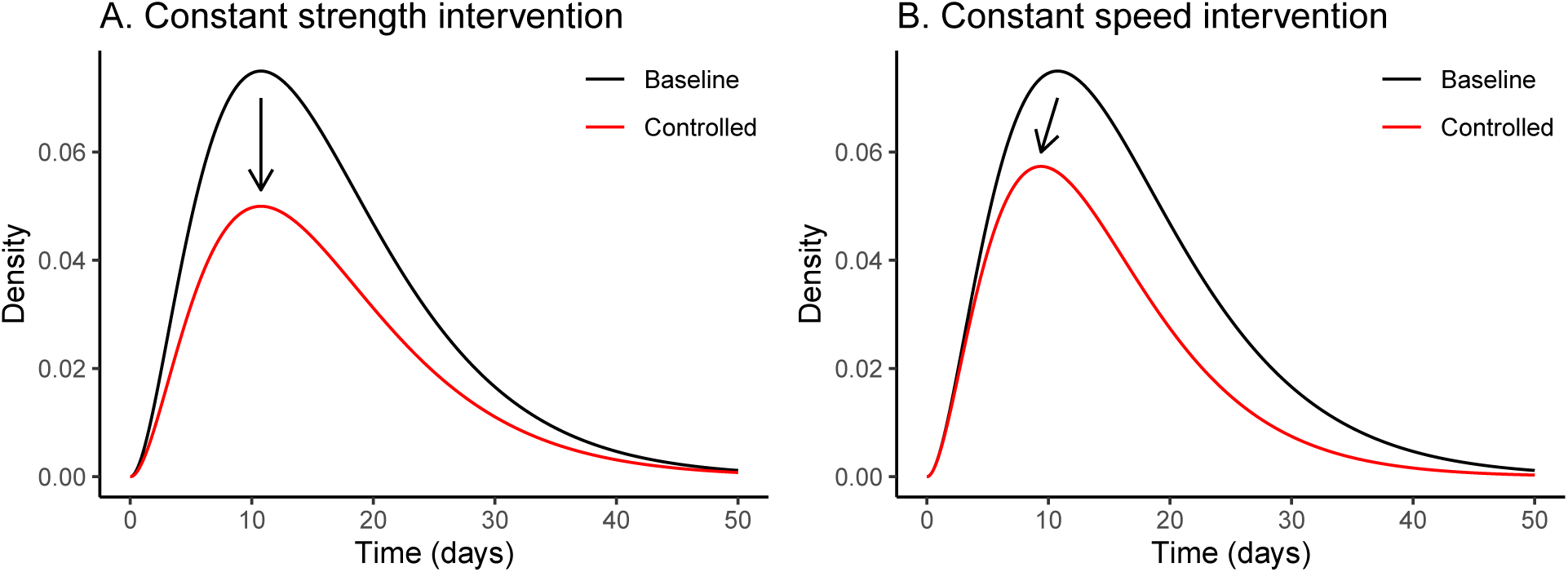
Effects of constant-strength and constant-speed intervention on infection kernels. Ebola-like gamma infection kernel *K*(*τ*) (mean: 16.2 days, CV: 0.58, and ℛ_0_: 1.5) is shown in black (Park et al., 2019). The infection kernel after applying each intervention strategy 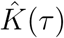 is shown in red. (A) The effect of a constant-strength intervention with *θ* = 1.5. A constant-strength intervention reduces the density by a constant proportion: 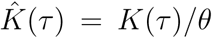; when the strength of intervention matches the strength of epidemic (*θ* = ℛ), the resulting distribution is equivalent to the intrinsic generation-interval distribution 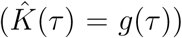. (B) A constant-speed intervention with *ϕ* ≈ 0.0267/day is applied to the same kernel. A constant-speed intervention reduces the density exponentially: 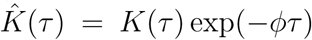; when the speed of intervention matches the speed of epidemic (*ϕ* = *r*), the resulting distribution is equivalent to the initial backward generation-interval distribution 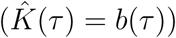.

Since *g* is a distribution, the reduction needed to prevent invasion (or to eliminate disease) is exactly *θ* = ℛ. We call *θ* the “strength” of the intervention; transmission is interrupted when the strength of the intervention *θ* is larger than the strength of spread ℛ.

We generalize this idea by allowing an intervention strategy to reduce *K* by different proportions over the course of an individual infection. We write the post-intervention kernel:

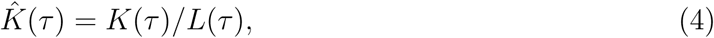

where *L*(*τ*) is the average proportional reduction for an individual infected time *τ* ago. The post-intervention reproductive number is thus:

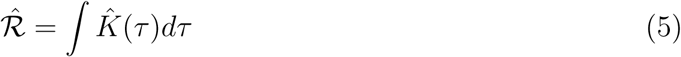

This framework generalizes the work of Fraser et al. (2004) who made parametric assumptions about the shape of *L*(*τ*).

We define the strength of the intervention *L* to be 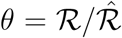. It is then straightforward to show that *θ* is the harmonic mean of *L*(*τ*) weighted by generation-interval distribution:

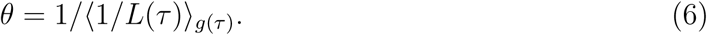

In this more general case, we have again that the disease cannot spread when *θ* ≥ ℛ.

### 2.3 Speed-based decomposition

The Euler-Lotka equation allows us to calculate the initial exponential growth rate *r* of an epidemic given an infection kernel *K*:

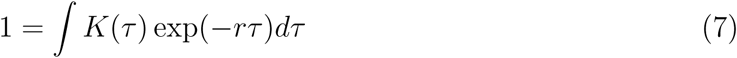

By analogy with the strength-based factorization (2), we can rewrite (7) as a speed-based factorization:

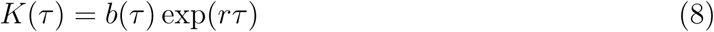

Like *g, b* is a distribution: in this case the initial backward generation interval, which gives the distribution of realized generation times (measured from the infectee’s point of view) when the disease spreads exponentially (Champredon and Dushoff, 2015; Britton and Scalia Tomba, 2019).

Now imagine an intervention that reduces transmission at a constant hazard rate *ϕ* across the disease generation (Fig. 1B), for example, by identifying and isolating infectious individuals. We then have:

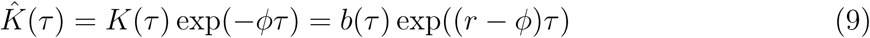

Since *b* is a distribution (which integrates to 1), the reduction needed to prevent invasion (or to eliminate disease) is exactly *ϕ* = *r*. We call *ϕ* the “speed” of the intervention; transmission is interrupted when the speed of the intervention is faster than the speed of spread.

We generalize this idea by allowing the hazard rate *h*(*τ*) at which *K* is reduced to vary through time, thus:

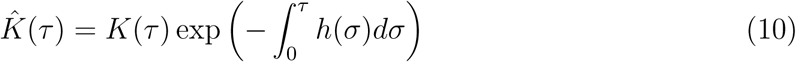

The associated post-intervention epidemic speed 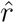 is given by:

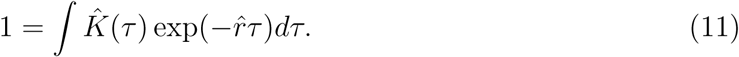

We define the speed of a general intervention to be 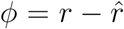. We can then show that *ϕ* is a (sort of) mean satisfying the implicit equation:

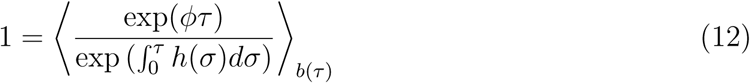

Specifically, the speed *ϕ* is a mean of the hazard *h* in the sense that an increase (or decrease) in *h* produces the same sign of change in *ϕ*, and if *h* is constant across the generation then *ϕ* = *h*.

## 3 Example: Human immunodeficiency virus (HIV)

In this section, we use both strength- and speed-based decompositions to compare different intervention strategies for the human immunodeficiency virus (HIV). In particular, we study how the amount of early HIV transmission affects estimates of intervention effectiveness. These examples are not detailed estimates for specific scenarios; instead, they are meant to demonstrate how strength- and speed-based decompositions can help evaluate control strategies.

We model the infection kernel of the HIV as a sum of two gamma distributions:

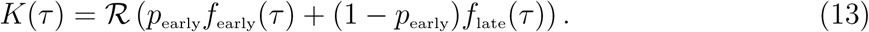

The first component, *f*_early_(*τ*), models early HIV transmission during the acute infection stage. We assume that *f*_early_(*τ*) has a mean of 3 months (Hollingsworth et al., 2008) and a shape parameter of 3. The second component, *f*_late_, models HIV transmission during the asymptomatic stage and the disease stage (after progression to Acquired Immune Deficiency Syndrome (AIDS)). We assume that *f*_late_(*τ*) has a mean of 10 years (Brookmeyer and Goedert, 1989; Nishiura, 2019) and a shape parameter of 2 (to roughly match the wide generation-interval distribution of HIV (Fraser et al., 2004)). Finally, *p*_early_ is the proportion of early HIV transmission.

The infection kernel is shown in (Fig. 2A) for our baseline value of *p*_early_ = 0.23. We assume that the initial speed of the epidemic is *r* = 0.452 year^−1^(Fig. 2B), and ask what value of ℛ_0_ would produce this rate of growth. When transmission is fast, (i.e., when *p*_early_ is large), individuals don’t need to transmit as much to achieve this speed, so the estimated value of ℛ_0_ decreases (Fig. 2C). Therefore, as *p*_early_ gets smaller, we expect stronger intervention to be required in order to control the disease.

**Figure 2:**
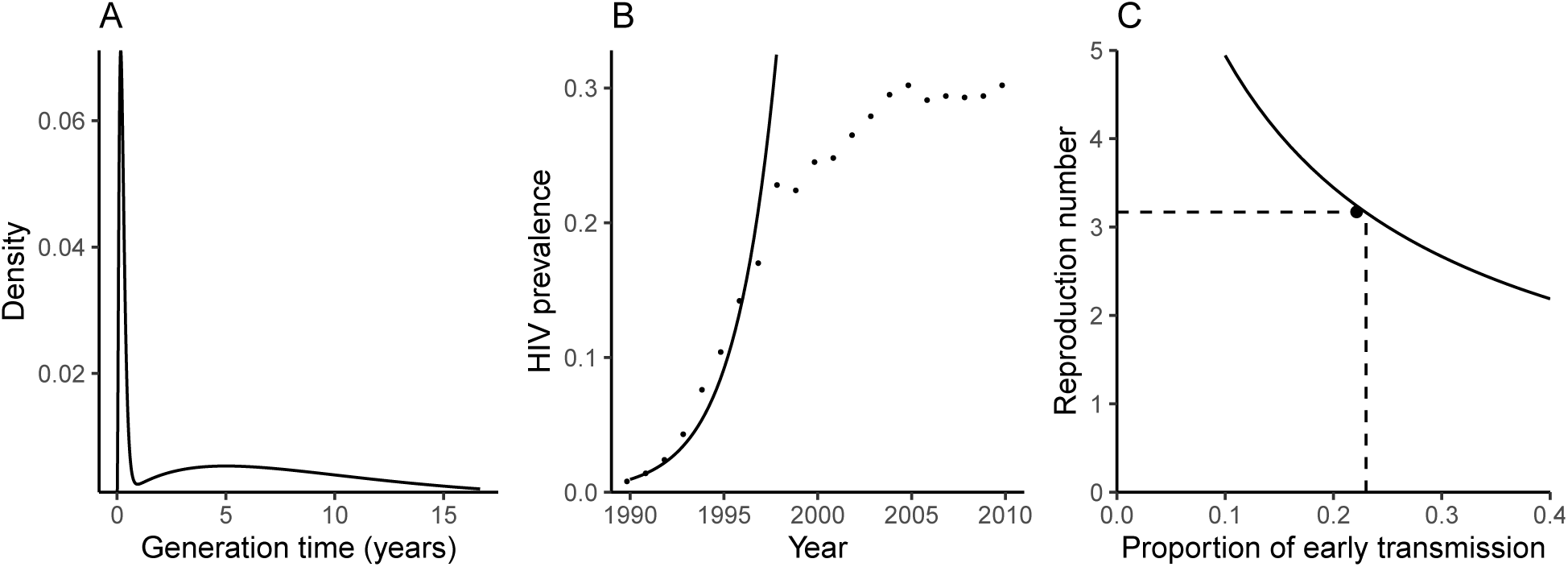
The infection kernel of the HIV. (A) The infection kernel of the HIV is approximated using a sum of two gamma distributions. We assume that the baseline proportion of early transmission is 23% (Hayes and White, 2006). (B) Time series of HIV prevalence in pregnant women in South Africa, 1990 - 2010 (Barron et al., 2013). The initial exponential growth rate of the HIV is estimated by fitting a straight line to log-prevalence (1990 - 1997) by minimizing the sum of squares. (C) Increase in the estimate of the amount of early transmission reduces the estimate of the reproductive number. The black circle indicates the baseline scenario.

We compare two different possible intervention strategies to shed light on the speed and strength decompositions. First, we consider a condom intervention that reduces HIV transmission by approximately 75% at the population level. Assuming that condoms act as a physical barrier, and that condom use will, on average, remain roughly constant through time, it is reasonable to model the proportional reduction in transmission due to condom use as constant across the course of infection: *L*_condom_ = 1/(1 − 0.75) = 4 (Fig. 3A). The estimated strength of such an intervention is simply the average of *L*_condom_, i.e., *θ* = 4, whereas the estimated strength of the epidemic decreases as the proportion of early transmission increases (Fig. 3B). Thus, the predicted effectiveness of the condom intervention will depend strongly on our estimate of the importance of early transmission: if early transmission is low, we expect disease spread to be too strong to be controlled completely by our intervention.

**Figure 3:**
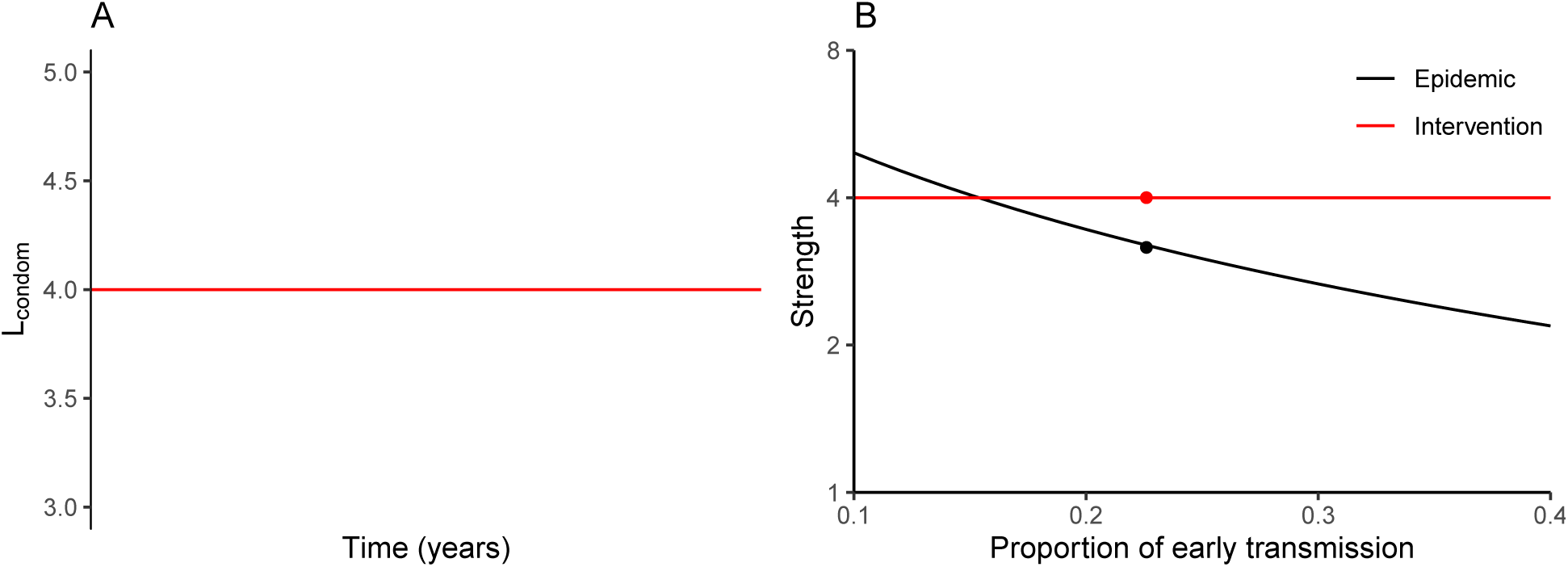
Evaluating a condom intervention using strength-based decomposition. (A) Condom use is thought to reduce probability of transmission by a similar factor throughout the course of infection; thus the proportional reduction *L*_condom_ due to condom use is constant across the course of infection. (B) The estimated amount of early transmission affects estimated strength of the epidemic but not of a condom-based intervention. The black and red circles indicate the baseline scenario.

Next, we consider a “test-and-treat” strategy in which infected individuals are identified, linked to care and receive antiretroviral therapy (ART) with the goal of both preserving health and preventing transmission through viral suppression. (Garnett and Baggaley, 2009; Granich et al., 2009; Nah et al., 2017). Our assumptions for this scenario are shown in Fig. 4. We assume that the hazard rate *h*_test_ of this intervention starts at 0 (because there is no way for newly infected individuals to know that they have HIV) but increases very quickly (because sexually active individuals are the most likely to seek testing); after a few months, the assumed hazard rate goes down to account for the effects of people who avoid identification, persistent treatment failures, and the possibility of rare transmission even under effective treatment (Fig. 4C). The corresponding strength of intervention *L* is shown in Fig. 4A and details of the assumption are given in the caption.

**Figure 4:**
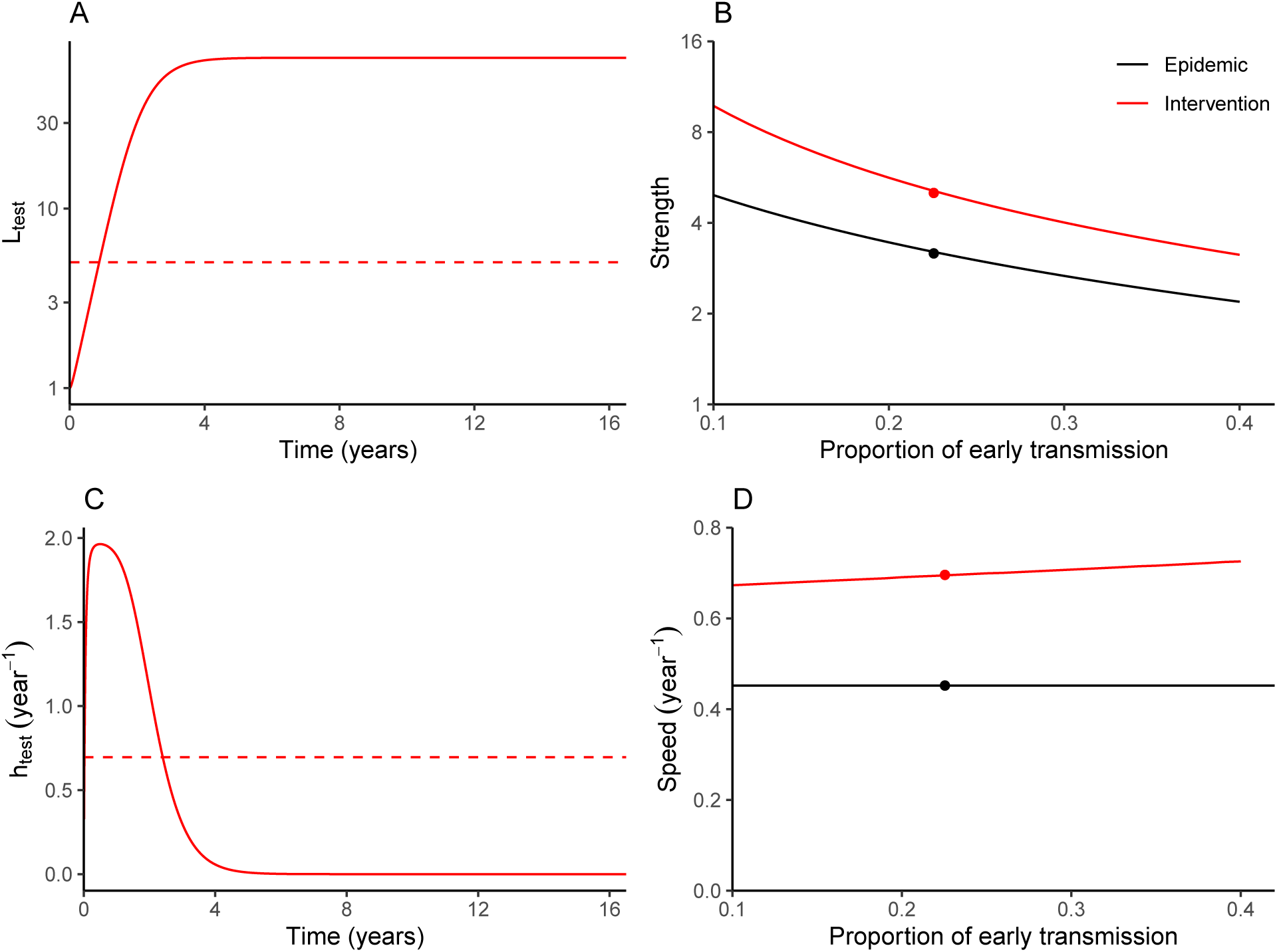
Evaluating a test-and-treat intervention using strength- and speed-based decomposition. (A) The strength of the test-and-treat intervention (calculated from the assumed hazard, (C)). The dashed line shows the corresponding effective strength of the intervention (from (6)) assuming 23% early transmission. (B) Increase in the estimated amount of early transmission decreases the estimated strength of an epidemic as well as the estimated strength of test-and-treat intervention. (C) The assumed hazard for the test-and-treat intervention. The dashed line shows the corresponding effective speed of the intervention (from (12)) assuming 23% early transmission. (D) The estimated amount of early transmission has little effect on the effective speed of intervention, and none on the speed of the epidemic estimated from incidence data. Circles indicate the baseline scenario. Test-and-treat intervention is modeled phenomenologically: 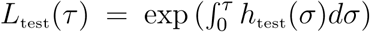 and *h*_test_(*τ*) = *h*_max_(1 − exp(−*Kf*(*τ*))), where *f*(*τ*) is a gamma probability density function with a mean of 1 year and a shape parameter of 2, *K* = 4/ max(*f*(*τ*)), and *h*_max_ = 2 year^−1^.

In this example, we see that, as *p*_early_ goes down and our estimate of epidemic strength increases, the estimate of intervention strength increases roughly in parallel. The increase in intervention strength makes sense, as we have more time to reach people on average before they transmit: this is the core of the result of Eaton and Hallett (2014). In our scenario, we predict that the intervention remains effective over the range of considered parameters.

Though there is a clear intuition for why both strengths increase as early transmission goes down, the speed paradigm provides insight into why these two increases are so close to parallel. The estimated epidemic speed depends only on the observed growth rate – it does not change if we change our assumption about the proportion of early transmission. For the test-and-treat intervention, the effective epidemic speed also stays relatively constant (Fig. 4D), in part because we have (plausibly) assumed that hazard stays relatively constant for a few key months, and in part because the backward generation-interval distributions for different scenarios are relatively similar. The intervention speed increases slightly as proportion of early transmission increases because the subpopulation that the intervention fails to reach become relatively more important if late transmission is more important. Thus, the speed paradigm provides an intuitive underpinning for the originally surprising result of Eaton and Hallett (2014): the effectiveness of test-and-treat interventions should not depend much on the proportion of early transmission.

## 4 Discussion

The effectiveness of an epidemic intervention is often measured by its ability to reduce the reproductive number – ℛ, or outbreak “strength” – below 1. The exponential growth rate – *r*, or outbreak “speed” – is often seen just as a stepping stone to ℛ or even overlooked entirely (Park et al., 2020). We argue that ℛ and *r* provide equally valid, complementary perspectives on epidemic control, and that there are situations where each provides a clearer picture than the other.

In this study, we: first extended the standard paradigm of ℛ as critical parameter for control, by defining the strength of an intervention on the same scale as ℛ, the strength of the epidemic; then constructed a parallel interpretation which measures the speed of an intervention on the same scale as *r*, the speed of an epidemic. We thus showed that the standard paradigm for ℛ and control has a natural parallel interpretation in terms of *r*.

To illustrate this idea, we used simple assumptions to explore the effects of two HIV intervention strategies (condoms and test-and-treat), using both strength- and speed-based frameworks. In particular, we provided an alternative explanation for the result of Eaton and Hallett (2014) who used detailed mathematical modeling of HIV transmission to show that the amount of early transmission does not affect the effectiveness of the ART: we can control an outbreak if we can identify infected individuals and enroll them on ART faster than the *observed* rate at which new cases are generated, which does not depend on the estimates of the amount of early transmission. The original explanation of the result relied on a strength-based argument: increasing the amount of early transmission decreases the basic reproductive number, which negatively correlates with the outcome of the ART intervention (Eaton and Hallett, 2014).

While both speed- and strength-based frameworks can give the same conclusion about the outcome of an intervention, sometimes one provides a clearer understanding of a given measure. For example, we expect the speed-based framework to be clearer for characterizing newly invading pathogens: when an epidemic is growing exponentially, the reproductive number cannot be estimated with confidence (Weitz and Dushoff, 2015), especially when there is large uncertainty in the shape of the generation-interval distribution (Park et al., 2020). Conversely, we expect the strength-based framework to be clearer for evaluating established pathogens (based on the effective proportion of the population susceptible). For interventions, we expect the speed-based framework to be clearer for evaluating intervention strategies that work at the individual level, like test-and-treat for HIV (Granich et al., 2009), or contact-tracing and quarantine for COVID-19 (Hellewell et al., 2020); we expect the strength-based framework to be clearer for intervention strategies that seek to reduce the overall transmission at the population level, like condom use. In other cases, such as real-time rollout of vaccines during an outbreak, both speed and strength approaches might be similarly uncertain because the result depends both on the speed of the rollout and the (strength-like) final coverage (Shah et al., 2018).

When comparing interventions with epidemic parameters to evaluate strategies, the situation is similar. Some scenarios lend themselves naturally to a single approach. For example, in the classic case of vaccination to eliminate a previously established childhood disease, both disease spread and intervention can be clearly characterized using strength (Anderson and May, 1985). In our HIV example, both the HIV epidemic and the test-and-treat intervention can be best characterized using speed. Other cases, such as using social distancing (a strength-like intervention) in the early stages of COVID-19 (epidemic speed is observed) may not fit so neatly into either paradigm, however.

There is an analogy here with measures of fitness in theoretical ecology. For example, when a population is regulated by density dependence that affects all individuals identically, *r* may be the best measure of fitness (Pasztor et al., 1996), but when regulation primarily affects juveniles mortality, ℛ is likely to be superior (Mylius and Diekmann, 1995). The importance of speed-based perspectives are still rarely recognized in the case of infectious disease, however.

Responses to the 2014 Ebola Outbreak in West Africa and the recent COVID-19 outbreak show an over-emphasis on strength at the expense of speed: during the early phases of both outbreaks, many disease modelers tried to estimate ℛ_0_ but overlooked *r*. For example, only 1 out of 7 preliminary analyses of the COVID-19 outbreak that were published as preprints between January 23–26, 2020 reported the doubling time of an epidemic (Bedford et al., 2020; Imai et al., 2020; Liu et al., 2020; Majumder and Mandl, 2020; Read et al., 2020; Riou and Althaus, 2020; Zhao et al., 2020). We suggest that infectious disease modelers should be aware of the complementarity of these two frameworks when analyzing disease outbreaks.

